# A Chromosome-level Assembly of the Japanese Eel Genome, Insights into Gene Duplication and Chromosomal Reorganization

**DOI:** 10.1101/2022.06.28.497880

**Authors:** Hongbo Wang, Hin Ting Wan, Bin Wu, Jianbo Jian, Alice HM Ng, Claire Yik-Lok Chung, Eugene Yui-Ching Chow, Jizhou Zhang, Anderson OL Wong, Keng Po Lai, Ting Fung Chan, Eric Lu Zhang, Chris Kong-Chu Wong

## Abstract

Japanese eels (*Anguilla japonica*) are commercially important species that have been harvested extensively for foods. Currently, this and related species (American and European eels) are difficult to breed on a commercial basis. Wild stock is used for aquaculture. Due to pollution, overfishing, and international trafficking, eel populations are declining. The International Union for Conservation of Nature lists Japanese eels as critically endangered and on its red list. Here we presented a high-quality genome assembly for Japanese eels and demonstrated that large chromosome reorganizations occurred in the events of third-round whole-genome duplications (3R-WRD). Following multiple chromosomal fusion and fission rearrangement, the Anguilla lineage has reduced the haploid chromosomal number of 19 from the ancestral proto-chromosomal number of 25. Phylogenetic analysis of expanded gene families showed the gene families of olfactory receptors and voltage-gated Ca^2+^-channel expanded significantly. The expansion of olfactory receptors (group δ and ζ genes) and voltage-gated Ca^2+^-channel gene families are important for olfaction and neurophysiological functions. Following 3R-WGD, additional tandem (TD) and proximal (PD) duplications occurred to acquire immune-related genes for adaptation. The Japanese eel assembly presented here can be used to study other Anguilla species that are related to evolution and conservation.

## INTRODUCTION

Fishes are highly diverse species living in many ecological habitats, including freshwater, estuarine, and the ocean^41^. Over 99% of fish species are known to be stenohaline, inhabiting either freshwater or marine environments. While euryhaline fishes are known to be diadromous, migrating between freshwater and marine environments in their life cycles^32^. Catadromous fishes like eels spawn in the sea and migrate to inland freshwater to grow and mature. Eels are ecologically and economically important, serving as indicators of the healthiness of coastal environments and resources in aquaculture. The fish are not bred in captivity^47^. In current practices, glass eels (juvenile life stage) are captured from the wild and raised on farms. Over 90 % of freshwater eels consumed worldwide are farm-raised. Since the 1960s, catches of Anguillid eels, like European and Japanese eels, have declined by over 50-80 %. In a 2014 report from the International Union for Conservation of Nature (IUCN), the American, European, and Japanese eels have been listed as being at the high risk of extinction. The decline in eel populations is abetted by soaring demand from global markets. In addition, overfishing, habitat loss, dams^75^, water pollution^26^, parasites^35^, eel larvae predation by mesopelagic fishes^48^, climate change, and altered ocean currents ^14^ are known to cause population decline.

From the evolutionary perspective, eels are among the most basal extant groups of teleost fishes and close to the non-teleost ray-finned fishes, including holostei (bowfin, gar), chondrostei (sturgeon, paddlefish, starlet), and cladistia (bichir, ropefish), those undergoing the first 2-round of vertebrate genome duplication, occurred before the divergence of ray fined and lobe-finned fishes 450 million years ago^42^. Eels are one of the first groups separated from the majority of the teleost fishes after the teleost-specific of whole-genome duplication (3R-WGD)^67^. The comparison of eels with other teleosts would shed light on fish evolution. In 2012, the first draft sequences of the genomes of Japanese (genome size 1.15 Gb, consisting of 323,776 scaffolds) and European eels (0.923 Gb, N50 of 78Kbp) were published^36,37^. The Japanese eel’s draft genome’s annotation was then enhanced using transcriptome data^60^. Moreover, the genome sequence assembly of the European eel (0.86 Gb genome size) was improved using nanopore sequencing^46^. A draft genome of the American eel (with a total size of 1.41 GB) was published in 2017, and 26,564 genes were annotated^73^. In 2019, the assembly of a Japanese genome of 1.13 Gb^17^ was improved with 256,649 contigs, 41,687 scaffolds, and a scaffold N50 of 1.03M. Currently, only the draft genome is available for Japanese eels. The purpose of this study was to provide high-quality genome assemblies and to understand the evolution of karyotypes in early ray-finned fishes. The genome-scale data can provide ecological and conservation information by identifying adaptive and disease-resistant alleles.

## RESULTS

### Genome Assembly and Annotation

In this study, MitoZ software was used to assemble and annotate the mitochondrial genome (16.686Kb) of our sample to confirm the species’ identity (**Methods**). The data matched with the Japanese eel mitochondrial genome (GenBank ID AB038556.2) of the NR database from NCBI (**Supplementary Fig 1&2**). We hierarchically integrated the sequencing data from different platforms to characterize their strength in *de novo* assembly and annotation (**Supplementary Fig. 3**). The draft genome was generated using ONT contigs followed by error correction and scaffolding based on the genomic spans of different sequencing technologies^28^ (**Methods**). A high-quality Japanese female eel’s reference genome was then generated, through the integration of ONT long reads (234x, 239.64Gb), PacBio CLR (261x, 267Gb), 10x Chromium linked-reads (313x, 319.7Gb), Hi-C data (48x, 48.99 Gb), Illumnia short-reads (148x, 151.89Gb) and mate-pair reads (127x, 130.5Gb). The contigs from ONT long reads resulted in a significantly improved N50 (25.82Mb) without losing many complete genes (54.6%) (**Supplementary Table 2**). With reduced assembly errors, the percentage of complete genes increased from 54.6% to 90.1%, indicating a higher base quality (**Supplementary Table 3**). For scaffolding, 10x linked-reads, Bionano, and Hi-C data were used sequentially according to fragment length to increase assembly continuity and assign scaffolds to 19 chromosomes (**Fig. 1, Supplementary Fig 3**, and **Supplementary Tables 4**). As a result, the genome size is 1.028Gb, the contig N50 is 21.48Mb, and the scaffold N50 is 58.7Mb. The chromosome lengths range from 19.93Mb to 94.28Mb. According to *actinopterygii_odb10* in the BUSCO database, 94% of the single-copy direct homologs in the Ray-finned Fishes were assembled in Japanese eels (**Supplementary Table 5**). The repeat elements were predicted to account for 30.49% of the whole genome (**Supplementary Tables 6)**. The TEs were excluded from gene annotation (**Supplementary Tables 7**). Japanese eels have a higher percentage (30.49%) of repetitive sequences, which may explain their larger genome, compared to European eels (*Anguilla anguilla* 0.979 Gb)^46^. Even so, the Japanese and European eels have a 1:1 correspondence pattern of chromosomes and 19,325 homologous genes, demonstrating their matching structure (**Supplementary Fig 4**).

**Figure 1.**
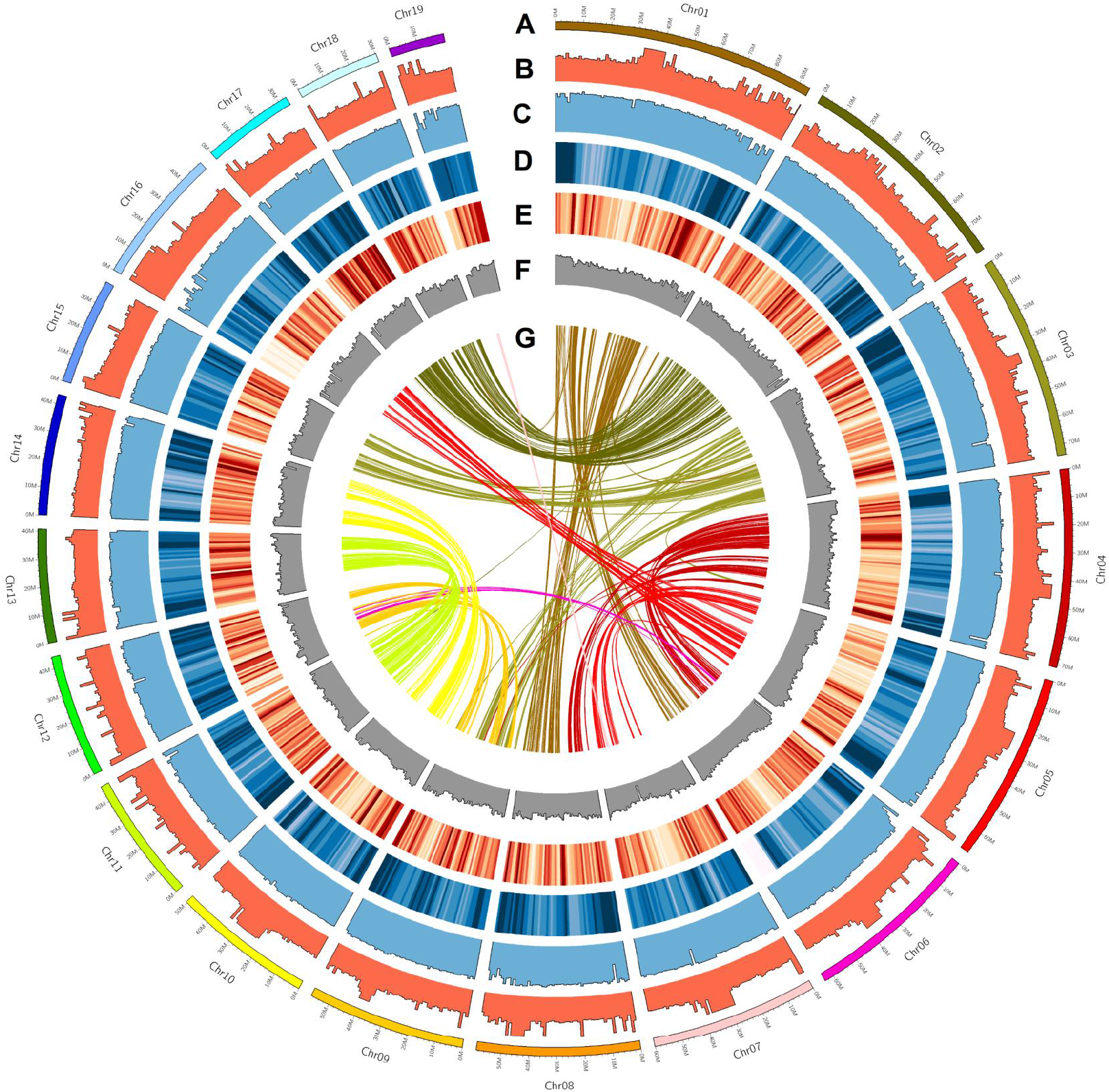
The genome landscape of Japanese eel, *Anguilla japonica*. From outer to inner circle: **(A)** length of 19 chromosomes (Mb); **(B)** Read-depth of ONT long-reads; **(C)** Read-depth of Illumina short-reads; **(D)** Distribution of transposon sequences; **(E)** Distribution of protein-coding gene; **(F)** GC content; **(G)** Collinear blocks of at least 10 genes in genome. The window size is 1MB.

By combining gene annotations from homology, *de novo*, and transcriptome annotations (**Methods**), we identified 29,982 coding genes (**Table 1**). We functionally annotated 97.44 % (29,219) of these genes (**Supplementary Table 8**) using the publicly available databases (**Methods**). Additionally, 21,606 genes were annotated by all five major protein databases (**Supplementary Fig 5)**, with signal transduction pathways most abundant in KEGG **(Supplementary Fig 6)** and KOG **(Supplementary Fig 7)**. BUSCO analysis showed that 94.7 % of the single-copy orthologs could be found in the ray-finned fish single-copy direct homology gene database actinopterygii_odb10 (**Supplementary Table 9**). The protein-coding genes in Japanese eels have an average length of 10.2kbp and contain approximately 9 exons (**Table 1**), which has an average length of 1.6kbp (**Supplementary Table 10**). The gene structure of Japanese eels is similar to those of four closely related species (**Supplementary Fig 8**). The genome assembly has a greater number of predicted genes (29,982 genes) than the Atlantic species, European (25,903 genes), and America (26,565 genes) eels. Additionally, 17,095 noncoding RNAs were predicted, including 1,042 transfer RNAs (tRNAs), 1,771 ribosomal RNAs (rRNAs), and 3,974 microRNAs in Japanese eels.

### Phylogenomics and demographic history

The orthology analysis of 12 species’ coding genes identified 21,653 gene family clusters. *Anguilla japonica*’s genome contains 29,982 coding genes, including 3347 single-copy orthologs, 8204 multiple-copy orthologs, 233 unique paralogs, 12,662 other orthologs, and 5536 unclustered genes. A phylogenetic tree was reconstructed by identifying the fourfold synonymous third-codon transversion (4dTv) loci in the 1,131 single-copy orthologs from the 12 fish species (**Fig 2**). American and European eels diverged from their ancestors about 27.0 million years ago (MYA). With a divergence time of approximately 44.1 MYA, Japanese eel was distant from the Atlantic eel species. The 4dTv-orthology density plot showed that Japanese eels are phylogenically closer to American than European eels. In comparison with the three freshwater eels (Anguilliformes) and tarpons (Elopiformes), the members of the Order Elopomorpha, their common ancestor, diverged 196.1 MYA. Elopomorpha and Osteoglossomorpha (i.e., arowana) are the closest evolutionary relatives at the basal branch of teleosts, separating 240.9 MYA. *Gadiformes* (e.g., Atlantic cod) and *Cypriniformes* (e.g., medaka, zebrafish) diverged from *Elopomorpha* and *Osteoglossomorpha* at 262.5 MYA. Above are groups of fishes, which had undergone three rounds of whole-genome duplication (3R-WGD). Compared to the outgroups, spotted gars, reed fish, coelacanths, and Australian ghost sharks underwent only 2R-WGD.

**Figure 2.**
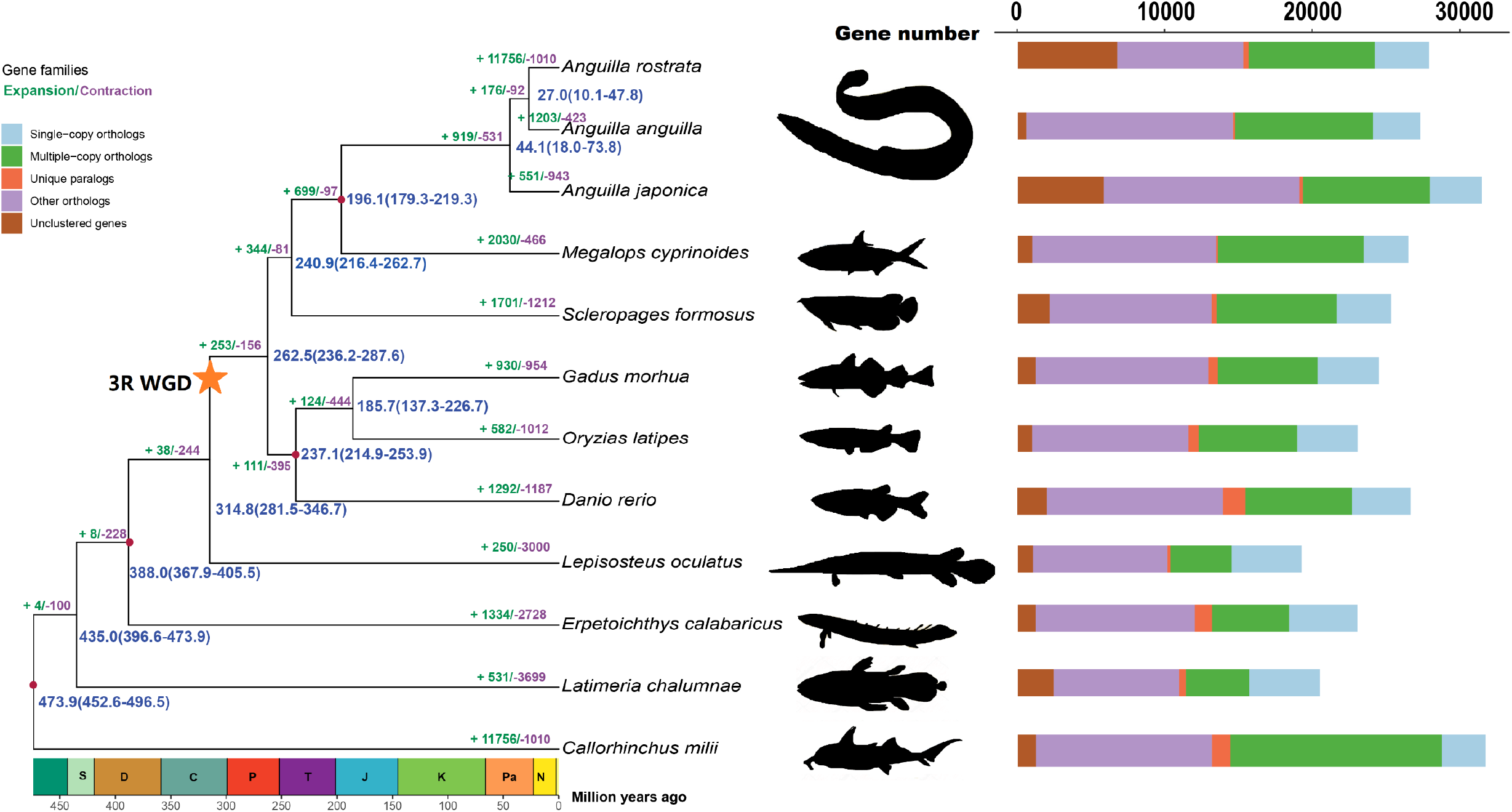
Phylogenetic relationship, divergence times and gene families of Anguilla species, relevant bony and cartilaginous fishes. Expansions (numbers in green) and contractions (numbers in purple) of gene families are shown at individual lineages. Each node shows the estimated divergence times (blue numbers, millions of years ago, Mya) and the 95% confidence intervals for these dates. Red dots indicate times taken from the TimeTree website (http://www.timetree.org/). The orange star shows the 3R-WGD event. Geological periods from left to right: S= Silurian, D= Devonian, C= Carboniferous, P= Permian, T= Triassic, J= Jurassic, K= Cretaceous, Pa=: Paleogene, N= Neogene. A comparison of gene families associated with orthologs and paralogs in Japanese eel and the 11 fish species.

### Expanded Gene Families and Gene duplication

The expansion and contraction of gene families reflect the evolution of organisms’ adaptations to their environments. Ortholog analysis of genes from the 12 species (**Methods**) identified 21,652 gene family clusters. By removing gene families with too many (≥ 200) or too few (≤ 2) genes, we achieved 129,862 genes to evaluate the expansion and contraction of gene families (**Fig 2**). Compared to the nine other species (**Methods**), the three freshwater eels had expanded 771 and contracted 467 gene families, resulting in an increase of 919 and loss of 531 genes, respectively (**Supplementary Table 11**). Among those, the three freshwater eel species exhibited a significant expansion in the olfactory receptor (OR) gene family, which is crucial for detecting odor molecules under varying environmental conditions. A retrospective analysis of the OR receptors across 10 species’ genomes was performed and seven types of OR receptors were identified [alpha (α), beta (β), gamma (γ), delta (δ), epsilon (ε), zeta (ζ) and eta (η)] based on a previous study^69^. Compared to other fish species, the Japanese eels had a significantly higher number of OR genes (394) (**Fig 3**), located on the four chromosomes - Chr4 (2 genes), Chr9 (153 genes), Chr11 (1 gene), and Chr12 (238 genes). Similarly, the European eel contains 392 OR genes. The δ and ζ genes are the major OR genes in the eels.

**Figure 3.**
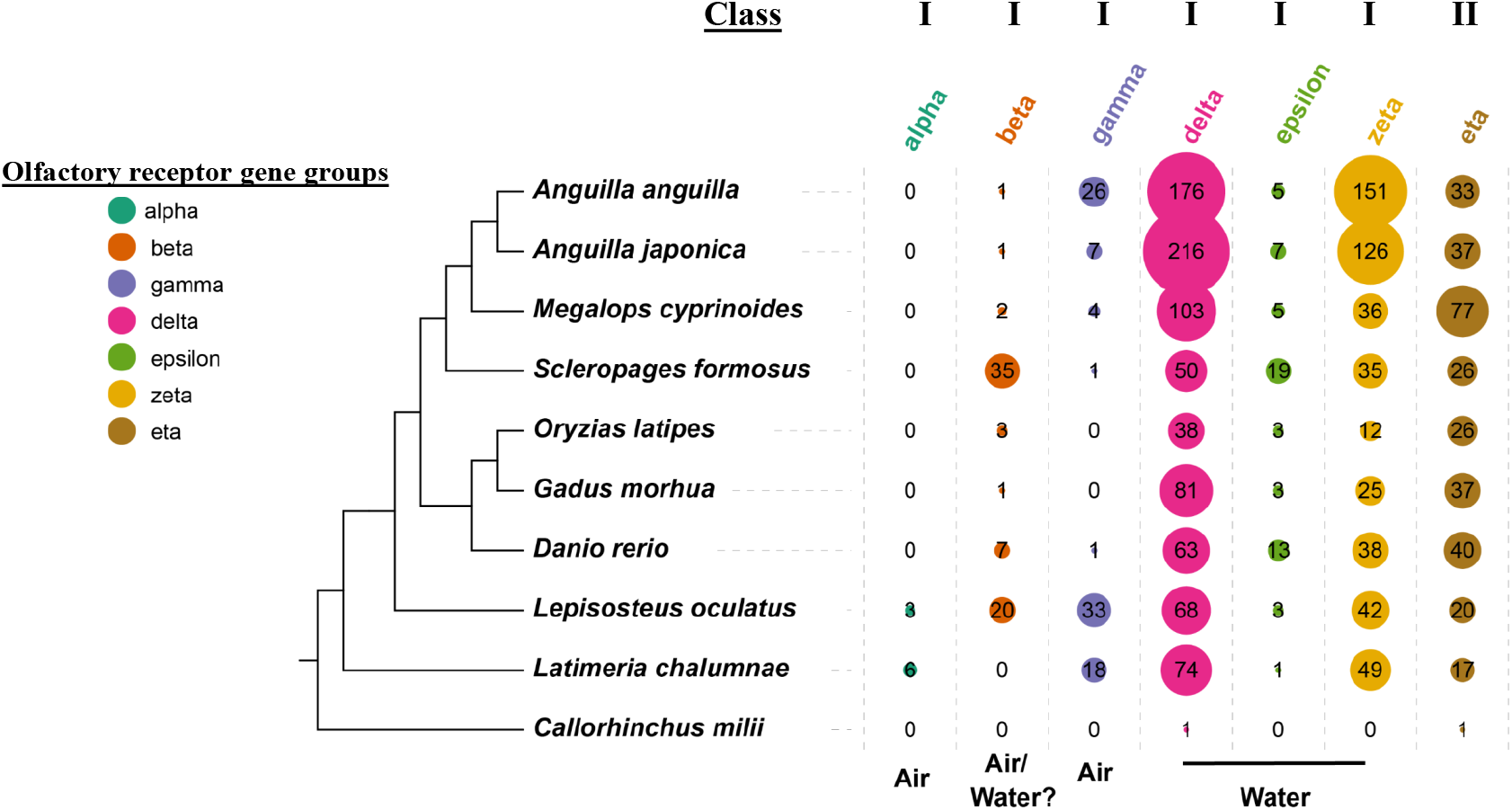
Number and classification of olfactory receptor (OR) genes for 10 fish species. On the left is the phylogenetic tree of the 10 species. The number of OR genes is shown on the right. The size of circle indicate the number of OR genes.

Comparing Japanese eel to the other 11 species, 433 gene families increased, with a total increase of 551 genes. A total of 943 genes were lost from 782 gene families (**Supplementary Table 12**). It is interesting to note that Ca^2+^ and K^+^ channel families were identified. Calcium and potassium play significant roles in neuronal excitability, muscle contraction, fertilization and energy metabolism. Interestingly, the other expanded gene families include (i) the assembly of myosin thick filament in skeletal muscle, (ii) lipoprotein receptor-related protein (metabolic and morphogenetic pathways) and (iii) isocitrate and isopropylmalate dehydrogenases family (carbohydrate and amino acid metabolism).

It was reported that freshwater eels (European and Japanese) had a large number of paralogous pairs, after splitting from the Osteoglossomorpha lineage^79^. The observation suggested 4R-WGD or lineage-specific rediploidization in some duplicated genomic regions. Our data showed that the genome size (1.028G) of Japanese eels is less than half that of the Atlantic salmon (2.97G)^59^, which underwent salmonid-specific 4R-WGD. In addition, we studied the distribution of 4dTv and Ks values of genome-wide direct homologous gene pairs in Japanese eels, European eels, and tarpons. There were 4dTv values of 0.402, 0.386, and 0.317 for *A. japonica, A. Anguilla, and M. cyprinoides*, respectively (**Fig 4A**, and **Supplementary Fig 9**). Additional WGD events were not detected. We also compared the syntenic blocks at Hox A-D loci with those in spotted gar that underwent 2R-WGD (**Fig 4B**). Japanese eel’s genome has eight clusters of Hox loci on chromosomes 1, 2, 3, 8, 11, 13, 15, and 17 while spotted gar has four clusters on chromosomes 4, 11, 12, and 13. Collectively, the data do not support the presence of 4R-WGD in Japanese and European eels.

**Figure 4.**
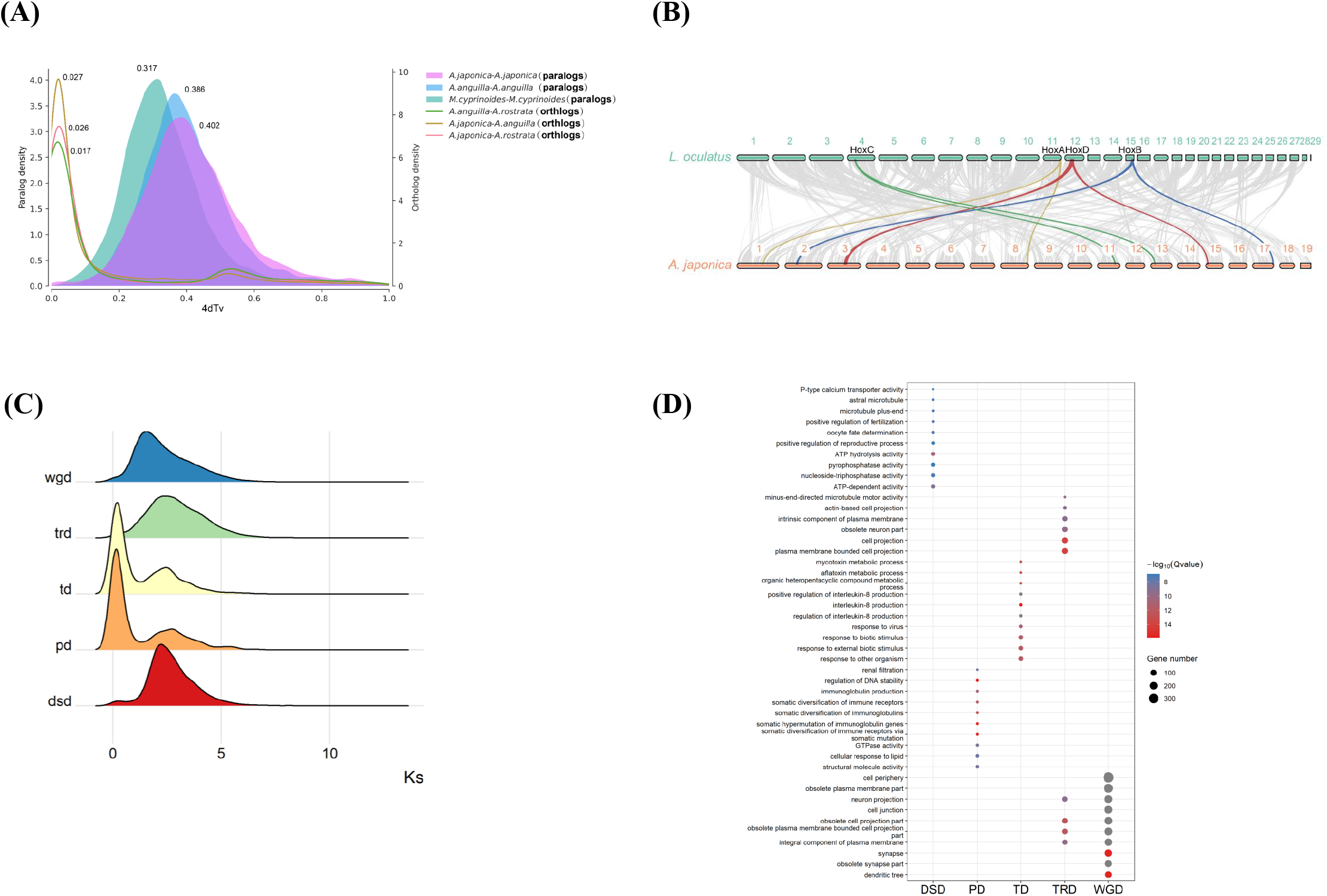
**(A)** Four-fold synonymous third-codon transversion rate (4dTv) distributions of homologous gene pairs for intra-species (paralogs density) and inter-species (orthologs density) comparisons. **(B)** The collinear relationships of syntenic blocks between *Anguilla japonica* and *Lepisosteus oculatus*. The numbers indicate the corresponding chromosomes for each species. In *Lepisosteus oculatus*, the 29th chromosome is 293.7 Kb long, which has no collinearity with that of *Anguilla japonica*. Based on homologous blocks of at least 10 genes, gene links between these two species were identified. The four collinear blocks that contain Hox genes are shown in green, yellow, red, and blue. **(C)** Ks distributions of syntenic gene pairs from different gene duplications (wgd: whole genome duplication, trd: transposable duplication, td: tandem duplication, pd: proximal duplication, dsd: dispersed duplication). The y-axis shows the distribution of Ks values. **(D)** Enrichment analysis of five duplicated expansion gene families, with the color of the circles representing the statistical significance of the GO. The circle size represents the number of genes.

There are a total of 21,249 duplicated genes identified among the 29,982 coding genes in the Japanese eel genome. Based on their duplication patterns, DupGen_finder (**Methods**) classified the duplicated genes into five categories, (i) 9,890 whole-genome duplicates (WGD, 46.54%), (ii) 1,420 tandem duplicates (TD, 6.68%), (iii) 768 proximal duplicates (PD, 3.61%), (iv) 3,975 transposed duplicates (TRD, 18.71%), and (v) 5,196 dispersed duplicates (DSD, 24.45%). We then calculated the Ks and Ka/Ks values for these five gene categories. Ks distribution indicates that TD and PD revealed additional duplication, post-3R-WGD **(Fig 4C)**. In addition, both TD and PD duplicates exhibited high Ka/Ks ratios, an indication of high selection pressure, which was probably related to environmental adaptation. TD and PD duplicated genes are mainly involved in immune responses (e.g. the production of interleukin-8, virus and biotic stress, somatic hypermutation of immunoglobulin genes, diversification and production of immunoglobulins and immunoreceptors) **(Fig 4D)**. Nonetheless, WGD was associated with 32.98 % of the total number of coding genes (29,982) in Japanese eels. Gene duplications in other fish species were also analyzed using the DupGen_finder pipeline (Qiao et al. 2019) and compared. Japanese eels were found to share the same level of WGD duplication of coding genes as arowana (37.60%), as both are extant members of the basal teleost group. However, it differs from the majority of teleosts, such as medaka (6.09%) zebrafish (9.51%) and Atlantic cod (4.68%). In Japanese eel, these duplicated gene functions were associated with neuronal (dendrites, synapses, neuron projections, obsolete synapses) and cell-cell junctions (cellular periphery, cell junctions, integral components of plasma membranes, obsolete plasma membranes, and cell projections). TRD shows a similar profile of changes. In DSD, duplication genes function in microtubules, reproduction (oocyte fate determination, fertilization), and ATP metabolism.

### Evolution of chromosome number in Japanese eels

When comparing chromosome numbers of the fishes all undergone 3R-WGD, the haploid chromosome number (n) is 25 for tarpons, arowana, zebrafish; 24 for medaka; 23 for Atlantic cod. Japanese eels have a lower haploid chromosome number (n = 19). To assess the extent of inter-chromosomal rearrangements in Japanese eels, we reconstructed the karyotype of the common ancestral teleosts karyotype (ATK) and ancestral eel-tarpon karyotype (AETK) (**Fig 5A**). According to our results, the ATK and AETK had 24 and 25 haploid chromosome numbers, respectively. The 14 AETK’s chromosomes (Chr1, 3, 4, 6, 7, 9, 16-20, 22, 23, 24) undergone 10-fusion and 10-fission to form the 14 tarpon’s chromosomes (Chr1, 4-8, 10, 11, 18, 19, 21-23, 25) (**Supplementary Table 13**). The remaining 11 AETK’s chromosomes (Chr2, 5, 8, 11, 25, 21, 15, 10, 12, 13, 14) correspond to those in tarpons (Chr2, 3, 9, 12-17, 20, 24). This chromosome rearrangement resulted in the same haploid chromosome number (n = 25) in tarpons. Comparatively, the 19 AETK’s chromosomes (Chr1, 2, 4, 7, 9-14, 16-21, 23-25) underwent 24-fusion and 18-fission to form the 13 chromosomes (Chr1-8, 11, 13, 15-17) in Japanese eels (**Supplementary Table 14**). The remaining 6 AETK’s chromosomes (Chr3, 5, 6, 8, 15, 22) correspond to the 6 chromosomes (Chr9, 10, 12, 14, 18 & 19) in Japanese eels. This chromosome rearrangement resulted in the reduction of the chromosome number (n = 19) in Japanese eels. Of which, Chr1, Chr3-7 rearrangements are unique to Japanese eels, and might play a role in speciation. Japanese eel’s chromosomes (Chr2, 11, 15) were essentially derived from AETK’s (Chr1, 3, 21) with slight rearrangements. The patterns of chromosome rearrangements in Chr8, Chr13, and Chr16-17 of Japanese eels were comparable with Chr21, Chr6, Chr8, and Chr19 in tarpons. Without rearrangement, Japanese eel’s chromosomes 10, 14 & 19 were equivalent to AETK’s chromosomes 3, 6, 22. In addition, there were 3 chromosomes in the Japanese eel (Chr9, Chr12, & Chr18) derived directly from AETK’s chromosomes (Chr5, Chr8, & Chr15), those also corresponding to tarpon’s chromosomes (Chr3, Chr9, & Chr15), respectively. **Figure 5B** and **Supplementary Fig 10** show the alignment of Japanese eel’s chromosomes to tarpon’s and arowana’s chromosomes and highlighted distinct conservation of orthologous segments.

**Figure 5.**
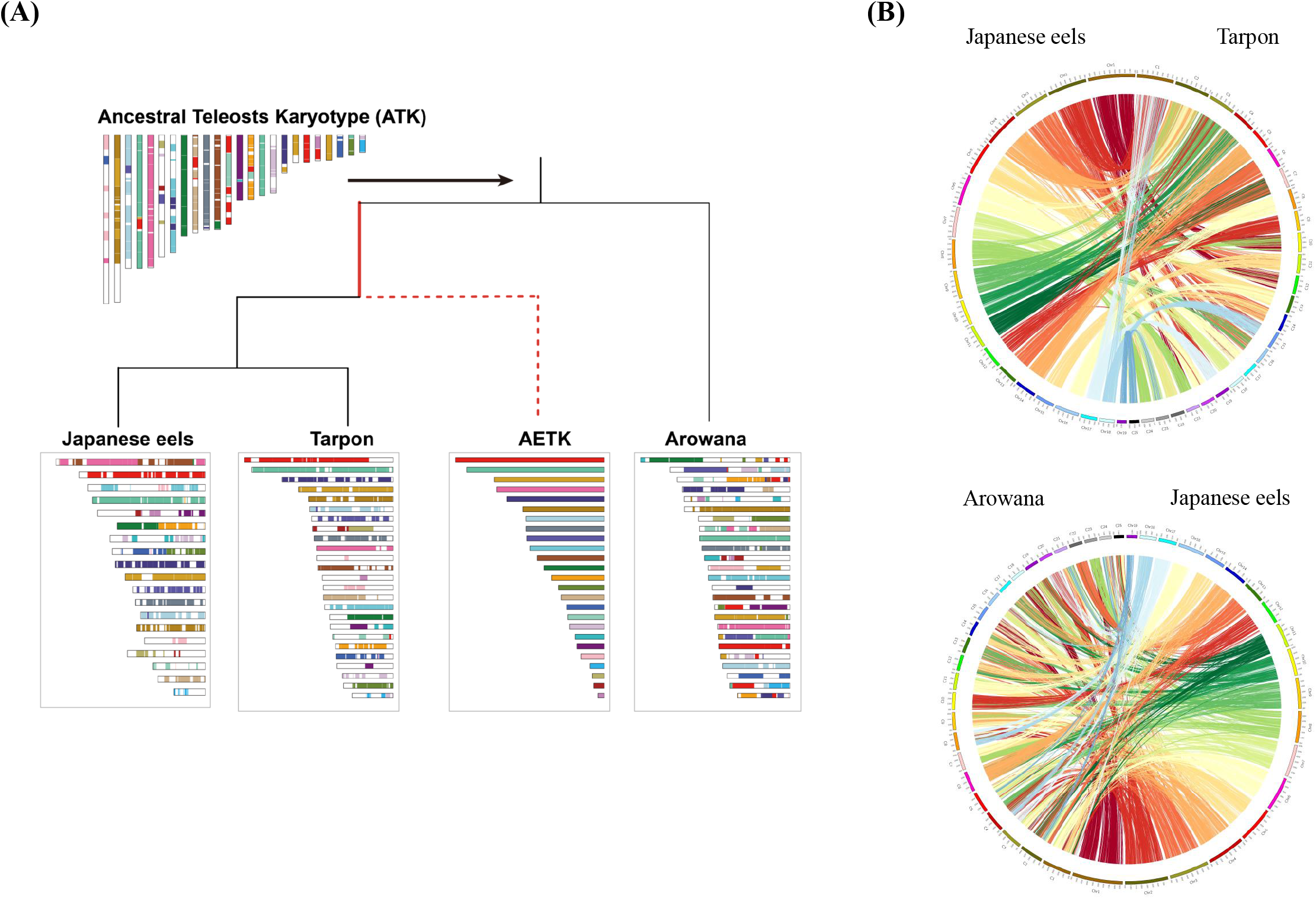
Reconstruction of proto-chromosomes for the common ancestor of teleosts (ATK) and eel/tarpons (AETK). **(A)** A model for the distribution of chromosomal segments in the genomes of ATK, arowana, AETK, Japanese eels and tarpons. AETK is the common ancestor of tarpons and eels. The Circos plots indicate conservation of synteny between **(B)** Japanese eel and tarpon, as well as **(C)** arowana and Japanese eel.

## DISCUSSION

In the past 10 years, the high-resolution whole-genome sequences of the teleosts, flatfish, zebrafish^40^, flatfish^16^, killifish^90^, salmon^59^, and the non-teleost ray-finned fishes, including spotted gar^11^, starlet sturgeon^22^, the early ray-finned fishes (i.e., bichir, paddlefish, bowfin & alligator gar)^7^ were published. However, as the basal extant group of teleosts, a high-resolution genome assemble of Atlantic or Pacific Anguilla species was not achieved. Here, we report the high-quality chromosomal-level Japanese eel’s genome for understanding the evolution of this basal extant group and providing the genome database for identifying adaptive and disease-resistant alleles.

The phylogenetic analysis of olfactory receptor (OR) genes identified from the genome sequences of medaka, Atlantic cod, zebrafish, gar, coelacanth and Australian ghost shark indicated that the delta (δ) and zeta (ζ) group genes in the freshwater eels expanded enormously, comprising about 86% of the entire gene family. Delta (δ) and ζ belong to the type I genes^7^, which are specialized for detecting water-soluble odorants and are uniquely expressed in the water-filled lateral diverticulum of the nasal cavity^25,30^. The mammalian type I (alpha group, α) and (gamma group, γ) genes detect airborne odor molecules. In teleost fishes, the group α genes are absent^7^. Interestingly, the group γ genes were found to have 26 in European eels and 7 in Japanese eels. Since eels can briefly live on land, they may have retained the group γ genes. The group β genes, which detect airborne and water-soluble odor molecules were low in the freshwater eels, but high in numbers in Arowana (35) and spotted gar (20). The group eta (η) genes (type 2) is the third major OR gene group in the freshwater eels. The group η genes are mainly expressed in fishes and is absent in mammals^70^.

The voltage-gated Ca^2+^ channels were found to be the major expanded gene families in Japanese eels. Genome studies suggest that the cellular functions of voltage-gated ion channels emerged early in Metazoan evolution^64,82^, in determining physiology and behavior at the time of early divergence. It is probably associated with the physiological challenge of Japanese eels to maintain a narrow range of intrinsic Ca^2+^, during migration between waters with great variations of calcium contents. A gene expression study in marbled eel (*Anguilla marmorata*) showed the high expression of voltage-gated Ca^2+^- channels in brain, skin and osmoregulatory tissues (i.e., gills, intestine, and kidneys), and its response to change in water calcium levels^12^. Besides controlling Ca^2+^ homeostasis, Ca^2+^-signaling coordinates a wide range of physiological processes, including skeletal muscle contractions, nervous system activity, cardiac and reproductive functions. The expanded gene families of myosin thick filament in skeletal muscle imply enhanced coordination of muscle contraction and performance^81^, especially for this distinct clade of elongated bodies inhabiting a diverse range of habitats^74^. Additionally, the expanded gene families in lipoprotein receptor-related protein, and the isocitrate and isopropylmalate dehydrogenases unravel the importance of this fundamental metabolic and morphogenetic functions in this lineage. Interestingly, lipoprotein receptor-related proteins first appeared during an evolutionary burst associated with the first multicellular organisms, and are multifunctional receptors in nervous system to modulate signals in brains^21,38^. Isocitrate dehydrogenase is an important enzyme of carbohydrate metabolism, while isopropylmalate dehydrogenase is involved in leucine biosynthesis. Although Japanese eels underwent 3R-WGD, an additional TD and PD duplication was detected. These duplication events, genetic raw materials were provided to facilitate new adaptations to the changing environment^65^. The duplicated gene functions of eels are immune-related to respond against different pathogens, which has been linked to the major factor contributing to the decline of eel populations^4,19,53^. Presumably, physiological fitness for adaptation might have been weakened by changes in the ecological environment, causing these evolutionary novelties^5^. Notably, the positive selection of immune-related genes indicates the adaptive advantages from the additional TD and PD duplication. Intriguingly, duplicated immune genes was also observed in salmon^55^ and sturgeon^22^.

The acquisition of evolutionary novelty by WGD duplication and the subsequent fate change of duplicated genes is necessary for phenotype alteration, environmental adaptation, and speciation^65^. The large scale of genomic reshaping after the third rounds of WGD affect evolutionary complexity and novelty in teleost fishes^29,43^. It has been widely established that chromosomal numbers are the most fundamental genomic characteristic of an organism or a lineage^62^. On the basis of the hypothesis that genome duplication resulted in chromosomal rearrangements^45^, an understanding of chromosome rearrangement in the eel genome may provide insight into the evolution of karyotype numbers at the base of the teleost evolutionary tree. The majority of fishes today have between 40 and 60 chromosomes (diploid number), while some commonly ancestral fishes are thought to have 48 chromosomes. Chromosome rearrangement and duplication have been the principal mechanisms involved in fish evolution, including the generation of new species and development of sex chromosomes. It is noted that freshwater fishes generally have higher number of chromosomes (the modal diploid number = 54) than marine fishes (the modal diploid number = 48). It has been suggested that the higher number of chromosomes in freshwater fishes is related to a less stable freshwater environment with greater topographical barriers^71^. A large capacity for dispersal in the marine environment, on the other hand, would contribute to homogenization of populations, reducing karyotype diversity^2^. Retrospectively, freshwater species seem to speciate more frequently than marine ones^9^. In a study of reconstructing the vertebrate ancestral genome to reveal dynamic genome reorganization, the 3R-WGD in the teleosts ancestor resulted in the number of chromosomes reaching haploid number (n) 26^66^. Evolutionarily, chromosome numbers peak at n=24 or 25 in extant teleost species. In this study, we were able to reconstruct the ancestral proto-chromosomes AETK (n = 25) to describe the cross-species chromosome collinearity and underpin the lineage-specific genome reorganization. The *Anguilla species* (n=19) diverged from *Megalops cyprinoides* (n=25) at 196.1 MYA, and their common ancestor from *Scleropages formosus* (n=25) at 240.9 MYA. The Anguilliformes are made up of 15 families with remarkable karyotypic diversity^91^. The haploid number ranges from 18 to 25, with a prevalence of n = 19 and 21. The *Anguilla* lineage underwent a significant structural rearrangement upon their divergence from tarpons (*Megalops cyprinoides*). The fusion and fission of their chromosome structure was the major drivers to reduce the haploid chromosome number to 19.

## METHODS

### Genome Sequencing

A market-purchased female Japanese eel, *Anguilla japonica* was kept in a freshwater tank for a week with aeration. Blood and muscle samples were taken from the fish, snapped frozen in liquid nitrogen, and then stored at -80°C. Genomic DNA was extracted from the blood samples. DNA sequencing data were generated by different platforms, including Oxford Nanopore (ONT) long reads, PacBio continuous long reads (CLR), Illumina short reads, Illumina mate-pair reads, 10X Chromium linked-reads, DNase Hi-C (Omni-C), and Bionano optical mapping.

For ONT long reads sequencing, the library was prepared by Ligation Sequencing Kit and sequenced using Nanopore PromethION P48 sequencer. For PacBio CLR sequencing, the SMRTbell templates were prepared using Sequel Binding Kit 1.0 and sequenced on the PacBio Sequel System. For Illumina short reads and mate-pair sequencing, the libraries were prepared using TruSeq DNA PCRFree Kit and Nextera Mate Pair Library Preparation Kit (gel plus), respectively. They were sequenced with 2×150bp reads on an Illumina HiSeq X Ten instrument. The library for linked-reads was prepared by 10X Genomics Chromium system with Chromium Genome library v2, and sequenced with 2×150bp reads on an Illumina NovaSeq 6000 instrument. Dovetail Omni-C Kit was used for Hi-C library preparation, which used NEBNext Ultra enzyme and Illumina-compatible adapters. Biotin-containing fragments were isolated using streptavidin beads prior to PCR enrichment.□The library was sequenced with 2×150bp reads on an Illumina HiSeqX platform. The Bionano optical mapping was generated by three enzymes, two from Irys (Nt.BspQI and Nb.BssSI) and one from Saphyr (DLE1). We stretched and captured the images of fluorescently labeled DNA molecules in Irys and Saphyr G1.2 chips. The labeling distances were extracted from the images and recorded into the raw molecule files. Molecules over 150 kbp were assembled into consensus maps using Bionano Solve for further analysis (**Supplementary Table 1**).

### Genome Assembly on ONT Long Reads

MitoZ software^63^ was used to assemble and annotate the mitochondrial genome of Japanese eel. We assembled ONT long reads using Canu^57^, Wtdbg2^80^, and Flye^56^ separately and merged their contigs using Quickmerge^13^ to achieve a balance between contig N50 and percentage of complete genes (PCGs) in vertebrate species. We used Racon^92^ for two rounds and Medaka (https://github.com/nanoporetech/medaka) for one round to self-correct assembly errors using ONT reads, respectively. The PacBio CLR were then incorporated for error correction using Racon for two rounds. As the last step, we further improved the assembly by integrating Illumina short-reads and mate-pair libraries using Pilon^93^ for two rounds.

### Scaffolding on 10x linked-reads, Bionano and Hi-C

We applied Tigmint^44^ and ARKS^18^ to correct misassembled contigs and linking contigs into scaffolds according to the shared barcodes from 10x linked-reads. We used OMGS^72^ to integrate three enzymes used in Bionano optical mapping for scaffolding. We further extended the scaffolds using 3D-DNA^23^ based on the Hi-C data from Dovetail Omni-C library and refined the scaffolds manually by JuiceBox^24^ to extend the scaffolds to the corresponding chromosome scale.

### Tandem Repeats and Transposable Elements Annotation

Tandem Repeats Finder v4.09^6^ was applied to annotate tandem repetitive sequences. We utilized homolog-based and *de novo* approaches to annotate transposable elements (TEs) in Japanese eel genome. For homolog-based approach, RepeatMasker v4.0.7^88^ and RepeatProteinMask v4.0.7 (http://www.repeatmasker.org/cgibin/RepeatProtein_MaskRequest) were used to identify the repeats by aligning the known TE sequences from RepBase v21.12 database^50^ to the genome. LTR_FINDER v1.07^96^ was used to infer long terminal repeat retrotransposons. For *de novo* approach, RepeatModeler (http://www.repeatmasker.org/RepeatModeler) was used to detect the TE families and repeat boundaries by integrating three complementary *de novo* repeat finding programs. RepeatMasker collected the union of these tools’ results and annotated the genome accordingly.

### Genes and their Functional Annotation

Three types of methods were used to annotate the protein-coding genes in the genome, including *de novo*, homology-based, and transcriptome-based annotations. Maker (v2.31.8)^39^ was adopted for homology annotation using the protein sequences from the five closely related species, including European eel (*Anguilla Anguilla)*, zebrafish (*Danio rerio*), Indo-Pacific tarpons (*Megalops cyprinoides*), Asian arowana (*Scleropages formosus*), and spotted gar (*Lepisosteus oculatus*), based on the phylogeny of teleost fishes^8^.

*De novo* annotation was performed using Augustus^85^ and SNAP^49^ by training a model using 3,000 complete genes obtained from homology prediction. Transcriptome annotation was performed by aligning RNA-seq data to the genome with HISAT2 (version 2.1.0)^54^, and assembling transcript sequences with Trinity^34^. Pasa_lite (https://github.com/PASApipeline/PASA_Lite) was used to correct assembly errors to obtain the final transcripts. Maker (v2.31.8) was further used to integrate the three annotations, followed by the second round of homology annotation to refine the final gene set.

Gene functional annotation was performed by aligning the predicted gene sequences to protein sequences using BLAST v2.2.31^1^ in the six databases, including NCBI Non-Redundant Protein Sequence (NR), Kyoto Encyclopedia of Genes and Genomes^51^, SwissProt^10^, KOG^89^, Gene Ontology^3^, and TrEMBL (Uniprot version 2020-06). We further searched the secondary structure domain database for gene function prediction using InterProscan^98^.

### Evaluation of Genome Assembly and Gene annotation

BUSCO^83^ was used to evaluate genome assembly and gene annotation by calculating the completeness of single-copy orthologs. We selected the Ray-finned Fish single-copy orthologs direct homologous gene database actinopterygii_odb10 (which contains 3640 core single-copy direct homologous gene proteins), the closest relative to Japanese eel in the OrthoDB database (https://www.orthodb.org/) to compare.

### Annotation of Conserved Noncoding Elements

tRNAscan-SE 1.3.1^61^ was used to identify tRNA sequences in the genome families. We annotated the rRNA sequences by aligning the conserved rRNA sequences from the five closely related fish species (European eel, zebrafish, tarpons, arowana, and spotted gar) to the genome using BLASTN^99^. The microRNAs and snRNAs were annotated by aligning the corresponding sequences from Rfam^31^ to the genome.

### Phylogenetic Analysis, Gene Expansion, and Gene Contraction

OrthoMCL 2.0^58^ (http://orthomcl.org/orthomcl/) was used to identify gene families by grouping orthologous proteins. We applied maximum likelihood method^33^ and RAxML^84^ (http://sco.h-its.org/exelixis/web/software/raxml/index.html) to reconstruct the phylogenetic tree using four-fold degenerate sites (4DTv) in single-copy orthologs from the 12 fish species, including *Anguilla rostrata* (American eel, GenBank assembly: GCA_001606085.1), *Anguilla anguilla* (European eel, GCA_013347855.1), *Anguilla japonica* (Japanese eel), *Megalops cyprinoides* (tarpons, GCA_013368585.1), *Scleropages formosus* (arowana, GCA_900964775.1), *Gadus morhua* (Atlantic cod, GCA_902167405.1), *Oryzias latipes* (medaka, GCA_002234675.1), *Danio rerio* (zebrafish, GCA_000002035.4), *Lepisosteus oculatus* (spotted gar, GCA_000242695.1), *Erpetoichthys calabaricus* (reed fish, GCA_900747795.2), *Latimeria chalumnae* (coelacanth, GCA_000225785.1) and *Callorhinchus milii* (Australian ghost shark, GCA_000165045.2). We estimated the divergence times for single-copy orthologos using mcmctree (http://abacus.gene.ucl.ac.uk/ software/paml.html) in PAML package^97^ based on the predefined times from TimeTree website (http://www.timetree.org/) [*Danio rerio* with *Oryzias latipes* (214.9 - 253.9 Mya), *Megalops cyprinoides* with *Anguilla anguilla* (179.3 - 219.3 Mya), *Callorhinchus milii* with *Danio rerio* (452.6 - 496.5 Mya) and *Erpetoichthys calabaricus* with *Danio rerio* (367.9 - 405.5 Mya)]. To estimate gene family expansion and contraction, we used CAFÉ^20^ (http://sourceforge.net/projects/cafehahnlab/) to model gene expansions and contractions, as well as the divergence times.

### Identification of olfactory receptor (OR) genes

We identified OR genes using the pipeline described in Github (https://github.com/MaximePolicarpo/Olfactory_receptor_genes)^76^, while candidate genes were filtered via the NR database. The OR gene identified in a previous study^69^ was used as a query sequence. TBLASTN^27^ was used to identify genomic regions containing OR genes in the 10 fish species (European eel, Japanese eel, tarpons, arowana, medaka, Atlantic cod, zebrafish, spotted gar, coelacanth, and Australian ghost shark). Only the non-overlapping BLAST hits regions were extracted, and the 1kb upstream and downstream flanking regions were used as the input to EMBOSS^78^. EMBOSS was used to generate the Open Reading Frames (ORFs), translated the ORFs into protein sequences, and then ran BlastP to weed out sequences that did not match genes already known in SwissProt and NR. InterProscan was used to determine the secondary structures of the predicted OR genes. Some genes were filtered due to lacking the seven transmembrane domains. The maximum likelihood phylogenetic tree was reconstructed using IQ-TREE^68^ based on the multiple sequencing alignments on the OR gene sequences with MAFFT^52^.

### Genome evolution analysis

MCscanX^95^ and macrosynteny visualization (jcvi) were used to screen for collinear blocks with at least 30 genes^87^ in *Anguilla japonica, Anguilla anguilla, Anguilla rostrata, Megalops cyprinoides, and Lepisosteus oculatus*. The numbers of non-synonymous substitutions (Ka) and synonymous substitutions (Ks) were calculated using KaKs_calculator2.0^94^. In addition, we calculated 4dTv values to estimate the WGD events in Japanese eel genome. We identified gene duplicates in the genomes of Japanese eel, zebrafish, arowana, medaka and Atlantic cod using the DupGen_finder pipeline^77^, using spotted gars as an outgroup. It classified gene duplication patterns into five categories: whole genome duplications, tandem duplications, proximal duplications (non-tandem duplications that are separated by 10 genes on the same chromosome), transposable duplications, and scattered duplications (duplications other than the four categories mentioned above).

### Ancestral Chromosome Reconfiguration

Ancestral eel/tarpon karyotype (AETK) was constructed using *Anguilla japonica* (Japanese eel), *Megalops cyprinoides* (tarpon) and *Scleropages formosus* (arowana, outgroup). The ancestral teleosts karyotype (ATK) was constructed using zebrafish, *Scleropages formosus* (arowana), and *Lepisosteus oculatus* (spotted gar, outgroup)^11^. This was done by using BLASTP^86^ to obtain homologous gene pairs between species. The default parameters of MCScanX were then used to obtain the collinear blocks of chromosomes between species. Finally, the karyotype of the ancestor was constructed using ANGeSv1.01^15^.

## Acknowledgement

This work was supported by the Southern Marine Science and Engineering Guangdong Laboratory (Guangzhou) (SMSEGL20SC02) to CKCW, AOLW, TFC & KPL, and General Research Fund (Research Grant Council, HKBU12162016) to CKCW.

## Data availability

The *Anguilla japonica* whole genome sequencing and assembly data has been deposited at National Library of Medicine, BioProject under the accession number PRJNA852364.

