## SupplementaryTables 1, 2, 3, 4, 5, 6, 7, 8, 9, 10, 13, 14 for "A Chromosome-level Assembly of the Japanese Eel Genome, Insights into Gene Duplication and Chromosomal Reorganization"

**Supplementary Table 1.** Genome sequencing platforms for *Anguilla japonica*.

| Reads | Reagents | Platforms |
| --- | --- | --- |
| Oxford Nanopore (ONT) long reads | Ligation Sequencing Kit | PromethION P48 |
| PacBio CLR long reads | Sequel Binding Kit 1.0 | PacBio Sequel |
| Illumina short reads | TruSeq DNA PCR Free Kit | Illumina HiSeq X Ten |
| Illumina mate-pair reads | Nextera mate pair library(gel plus) | Illumina HiSeq X Ten |
| DNAse Hi-C (Omni-C) reads | NEBNext Ultra enzyme | Illumina HiSeqX |
| Bionano optical mapping | Nt.BspQI, Nb.BssSI and DLE1 | Bionano Irys and Bionano Saphyr |
| 10X Chromium linked-reads | Chromium Genome library v2 | NovaSeq 6000 |

**Supplementary Table 2.** A summary of contig statistics from the ONT long read assembly.

| **Statistics** | **Flye** | **Wtdbg2** | **Canu** | **Quickmerge** |
| --- | --- | --- | --- | --- |
| Total contig lengths (Gb) | 1.056 | 1.051 | 1.476 | 1.061 |
| Contig N50 (Mb) | 4.6 | 11.49 | 3.99 | 25.82 |
| No. of contigs | 3,020 | 5,072 | 5,923 | 2,380 |
| Busco (complete) | 56.70% | 24.90% | 62.50% | 54.60% |

**Supplementary Table 3.** A summary of contig statistics after assembly error correction.

| **Statistics** | **Racon**  **(2 rounds)** | **Medaka** | **Racon (PacBio)** | **Pilon**  **(2 rounds)** |
| --- | --- | --- | --- | --- |
| Total contig lengths (Gb) | 1.07 | 1.08 | 1.08 | 1.07 |
| Contig N50 (Mb) | 26.10 | 19.79 | 19.79 | 19.66 |
| No. of contigs | 2,014 | 2,230 | 2,072 | 2,072 |
| Busco (complete) | 69.00% | 83.70% | 86.10% | 90.10% |

**Supplementary Table 4.** Scaffolding by 10x linked-reads, Bionano optical mapping and Hi-C.

| **Statistics** | **Tigment+ARKS** | **OMGS** | **3D-DNA** |
| --- | --- | --- | --- |
| Total contig lengths (Gb) | 1.07 | 1.10 | 1.028 |
| Contig N50 (Mb) | - | - | 21.48 |
| Scaffold N50 (Mb) | 21.53 | 25.37 | 58.70 |
| No. of contigs | 2,122 | 1,956 | 2,122 |
| Busco (complete) | 90.40% | 93.90% | 94.0% |

**Supplementary Table 5.** The completeness of *Anguilla japonica* genome by BUSCO assessment.

| **Complete BUSCOs** | **Complete and single-copy BUSCOs** | **Complete and duplicated BUSCOs** | **Fragmented BUSCOs** | **Missing BUSCOs** | **Total Lineage BUSCOs** |
| --- | --- | --- | --- | --- | --- |
| 3,423(94.0%) | 3,127(85.9%) | 296(8.1%) | 76(2.1%) | 141(3.9%) | 3,640(100%) |

**Supplementary Table 6.** Statistical results for repeat sequences.

| **Tools** | **Repeat Size** | **% of genome** |
| --- | --- | --- |
| Tandem Repeats Finder | 90,282,378 | 8.84 |
| RepeatMasker | 118,046,542 | 11.56 |
| RepeatProteinMask | 23,966,281 | 2.35 |
| RepeatModeler | 249,739,910 | 24.46 |
| Total | 311,306,842 | 30.49 |

**Supplementary Table 7.** A statistical analysis of the classification results for TE.

| **Types** | **TEs length** | **% in genome** | |
| --- | --- | --- | --- |
| DNA | 177.27Mb |  | 177.27Mb |
| LINE | 84.12Mb |  | 84.12Mb |
| SINE | 14.43Mb |  | 14.43Mb |
| LTR | 37.8Mb |  | 37.8Mb |
| Other | 42.79Kb |  | 42.79Kb |
| Unknown | 9.78Mb |  | 9.78Mb |
| Total | 264.02Mb |  | 264.02Mb |

**Supplementary Table 8.** Functional annotation of predicted genes from *Anguilla japonica*

| **Values** | **Total** | **NR-Annotated** | **Swissprot-Annotated** | **KEGG-Annotated** | **KOG-Annotated** | **TrEMBL-Annotated** | **Interpro-Annotated** | **GO-Annotated** | **Overall** |
| --- | --- | --- | --- | --- | --- | --- | --- | --- | --- |
| Number | 29,851 | 28,717 | 26,838 | 25,579 | 22,731 | 28,994 | 27,154 | 19,491 | 29,088 |
| Percentage | 1 | 0.962 | 0.8991 | 1 | 0.7615 | 0.9713 | 1 | 0.6529 | 0.9744 |

**Supplementary Table 9.** The completeness of *Anguilla japonica* genes by BUSCO assessment

| **Complete BUSCOs** | **Complete and single-copy BUSCOs** | **Complete and duplicated BUSCOs** | **Fragmented BUSCOs** | **Missing BUSCOs** | **Total Lineage BUSCOs** |
| --- | --- | --- | --- | --- | --- |
| 3446(94.7%) | 3136(86.2%) | 310(8.5%) | 50(1.4%) | 144(3.9%) | 3640(100%) |

**Supplementary Table 10.** The average length of exons in Japanese eel and the eight related fish species

| Species | Average exon length(bp) |
| --- | --- |
| *Anguilla japonica* | 1,600.77 |
| *Anguilla anguilla* | 1,827.74 |
| *Anguilla rostrata* | 1,516.78 |
| *Danio rerio* | 1,623.84 |
| *Lepisosteus oculatus* | 1,694.42 |
| *Megalops cyprinoides* | 1,796.75 |
| *Scleropages formosus* | 1,683.78 |
| *Gadus morhua* | 1,831.99 |
| *Erpetoichthys calabaricus* | 1,712.44 |

**Supplementary Table 13.** The karyotypes of *M. cyprinoides* (tarpons) and the common ancestor of eels and tarpons (AETK)

| *M. cyprinoides* | AETK |
| --- | --- |
| Chr01 | AETK_pChr1 |
| Chr02 | AETK_pChr2 |
| Chr03 | AETK_pChr5 |
| Chr04 | AETK_pChr3 |
| Chr05 | AETK_pChr6,AETK_pChr3,AETK_pChr6 |
| Chr06 | AETK_pChr7,AETK_pChr9,AETK_pChr7 |
| Chr07 | AETK_pChr9 |
| Chr08 | AETK_pChr23,AETK_pChr24 |
| Chr09 | AETK_pChr8 |
| Chr10 | AETK_pChr4 |
| Chr11 | AETK_pChr21 |
| Chr12 | AETK_pChr11 |
| Chr13 | AETK_pChr25 |
| Chr14 | AETK_pChr21 |
| Chr15 | AETK_pChr15 |
| Chr16 | AETK_pChr10 |
| Chr17 | AETK_pChr12 |
| Chr18 | AETK_pChr19,AETK_pChr20 |
| Chr19 | AETK_pChr1,AETK_pChr17 |
| Chr20 | AETK_pChr13 |
| Chr21 | AETK_pChr16,AETK_pChr4,AETK_pChr16 |
| Chr22 | AETK_pChr20 |
| Chr23 | AETK_pChr18 |
| Chr24 | AETK_pChr14 |
| Chr25 | AETK_pChr18,AETK_pChr22 |

**Supplementary Table 14.** The karyotypes of *A. japonica* (Japanese eel) and the common ancestor of eels and tarpons (AETK)

| ***A. japonica*** | **AETK** |
| --- | --- |
| Chr01 | AETK_pChr4,AETK_pChr12,AETK_pChr11,AETK_pChr4,AETK_pChr11,AETK_pChr4,AETK_pChr11 |
| Chr02 | AETK_pChr1 |
| Chr03 | AETK_pChr10,AETK_pChr21,AETK_pChr10 |
| Chr04 | AETK_pChr2,AETK_pChr13,AETK_pChr2 |
| Chr05 | AETK_pChr20,AETK_pChr25,AETK_pChr24 |
| Chr06 | AETK_pChr13,AETK_pChr12, |
| Chr07 | AETK_pChr19,AETK_pChr18,AETK_pChr19,AETK_pChr18 |
| Chr08 | AETK_pChr14,AETK_pChr16,AETK_pChr4,AETK_pChr16 |
| Chr09 | AETK_pChr5 |
| Chr10 | AETK_pChr3 |
| Chr11 | AETK_pChr9 |
| Chr12 | AETK_pChr8 |
| Chr13 | AETK_pChr7,AETK_pChr9,AETK_pChr7 |
| Chr14 | AETK_pChr6 |
| Chr15 | AETK_pChr21 |
| Chr16 | AETK_pChr24,AETK_pChr23 |
| Chr17 | AETK_pChr17,AETK_pChr1,AETK_pChr17 |
| Chr18 | AETK_pChr15 |
| Chr19 | AETK_pChr22 |
