## Supplementary Figures 1, 2, 3, 4, 5, 6, 7, 8, 9, 10 for "A Chromosome-level Assembly of the Japanese Eel Genome, Insights into Gene Duplication and Chromosomal Reorganization"

### Slide 1
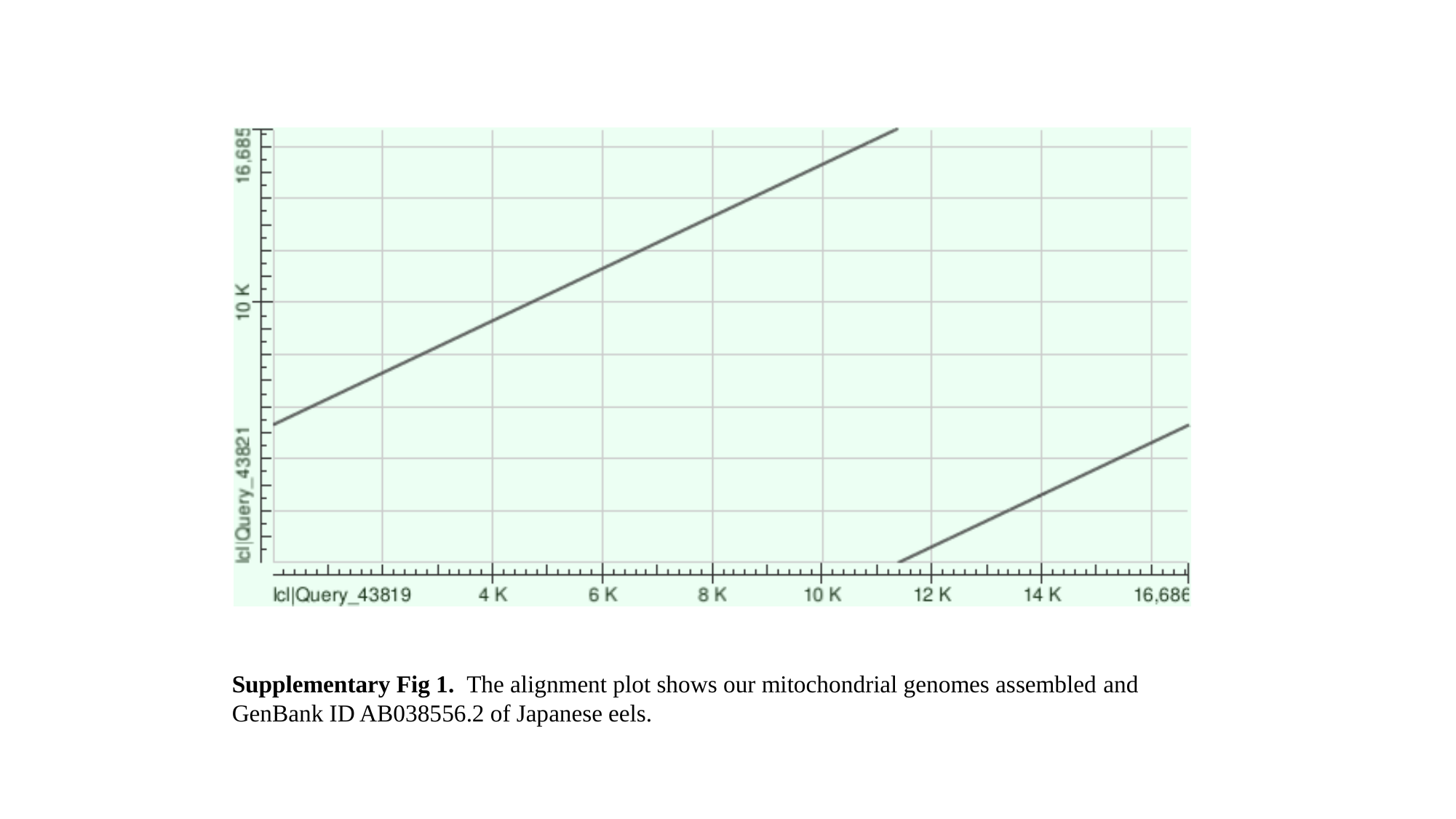

Supplementary Fig 1. The alignment plot shows our mitochondrial genomes assembled and GenBank ID AB038556.2 of Japanese eels.

### Slide 2
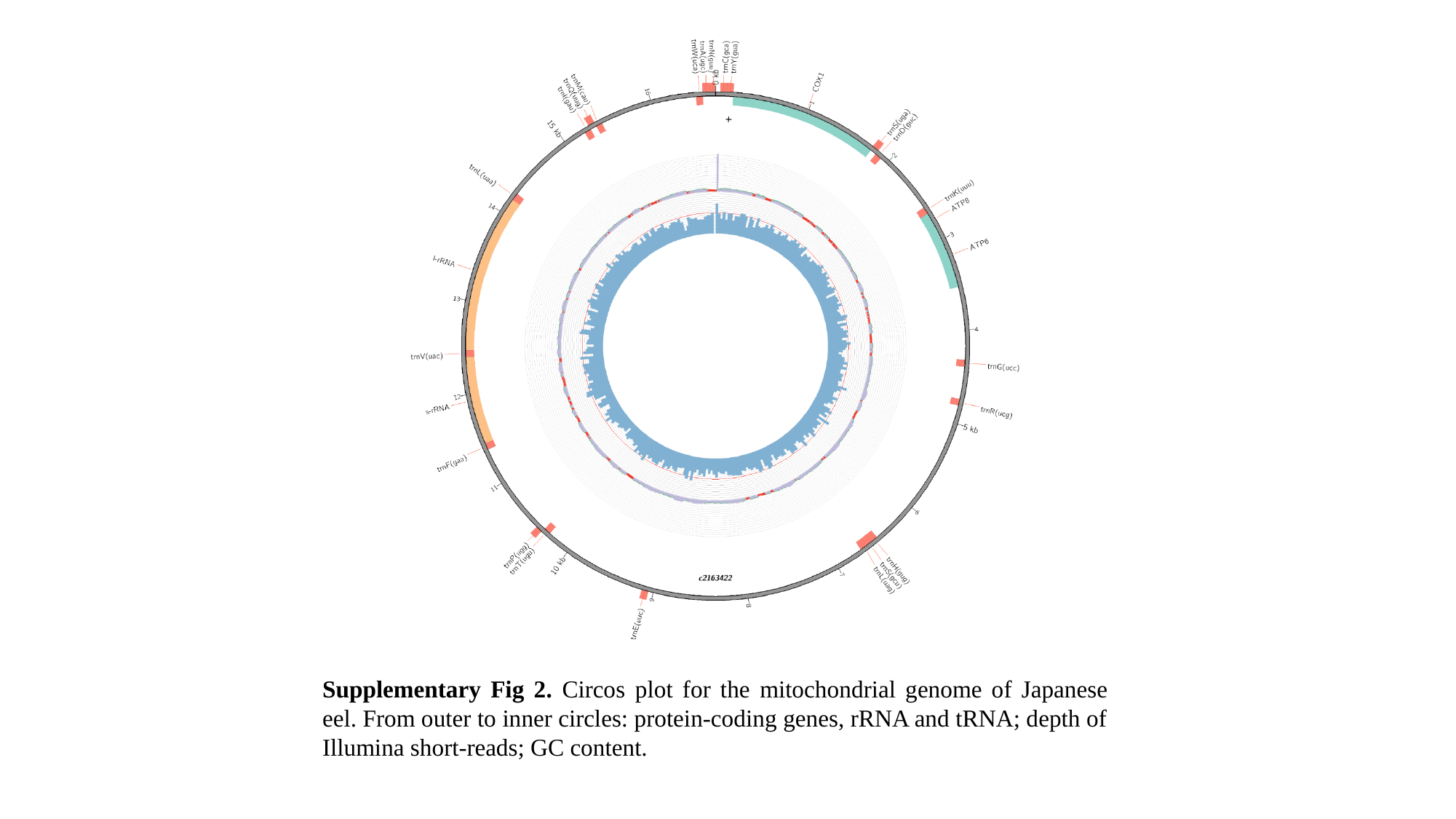

Supplementary Fig 2. Circos plot for the mitochondrial genome of Japanese eel. From outer to inner circles: protein-coding genes, rRNA and tRNA; depth of Illumina short-reads; GC content.

### Slide 3
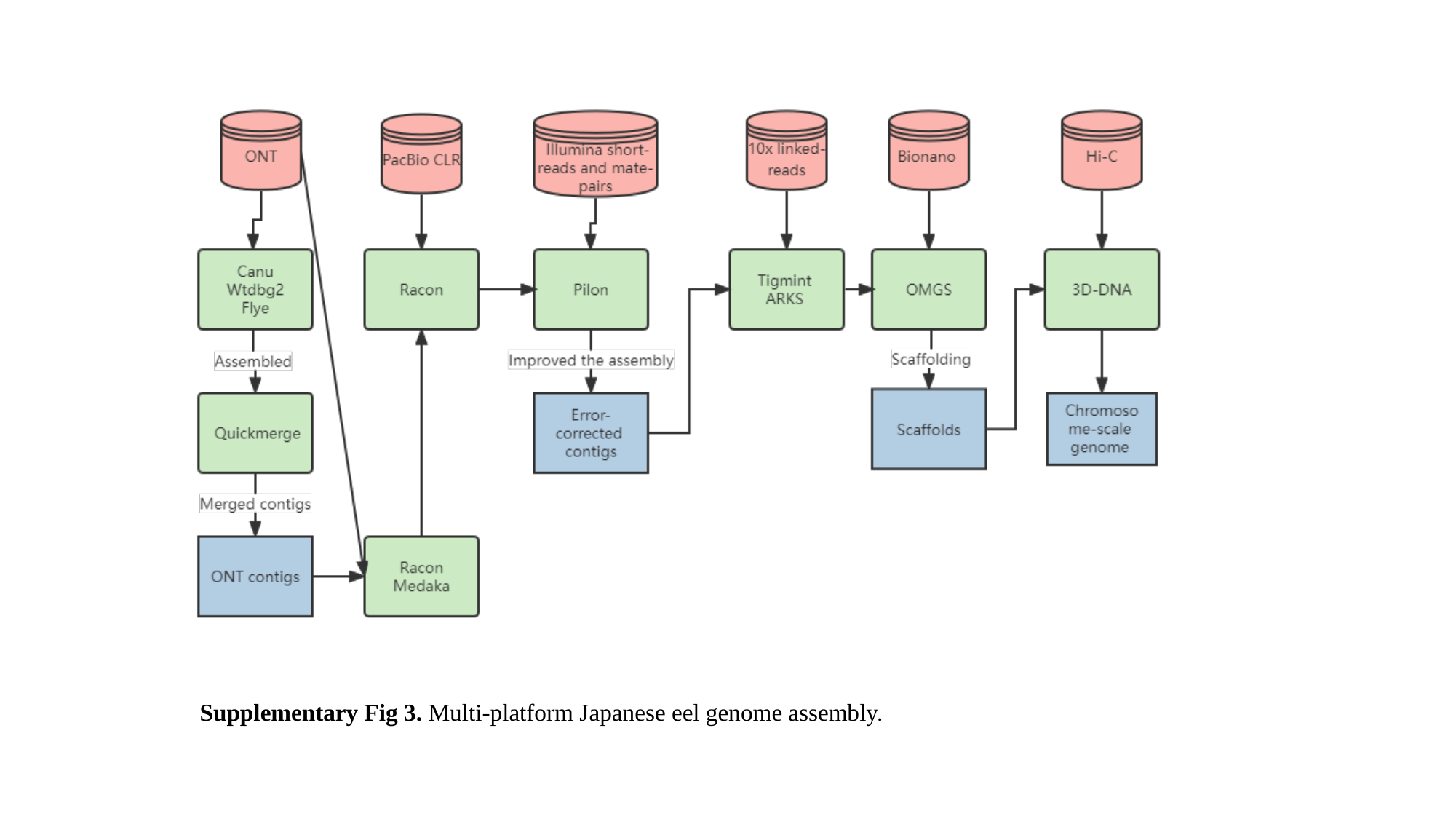

Supplementary Fig 3. Multi-platform Japanese eel genome assembly.

### Slide 4
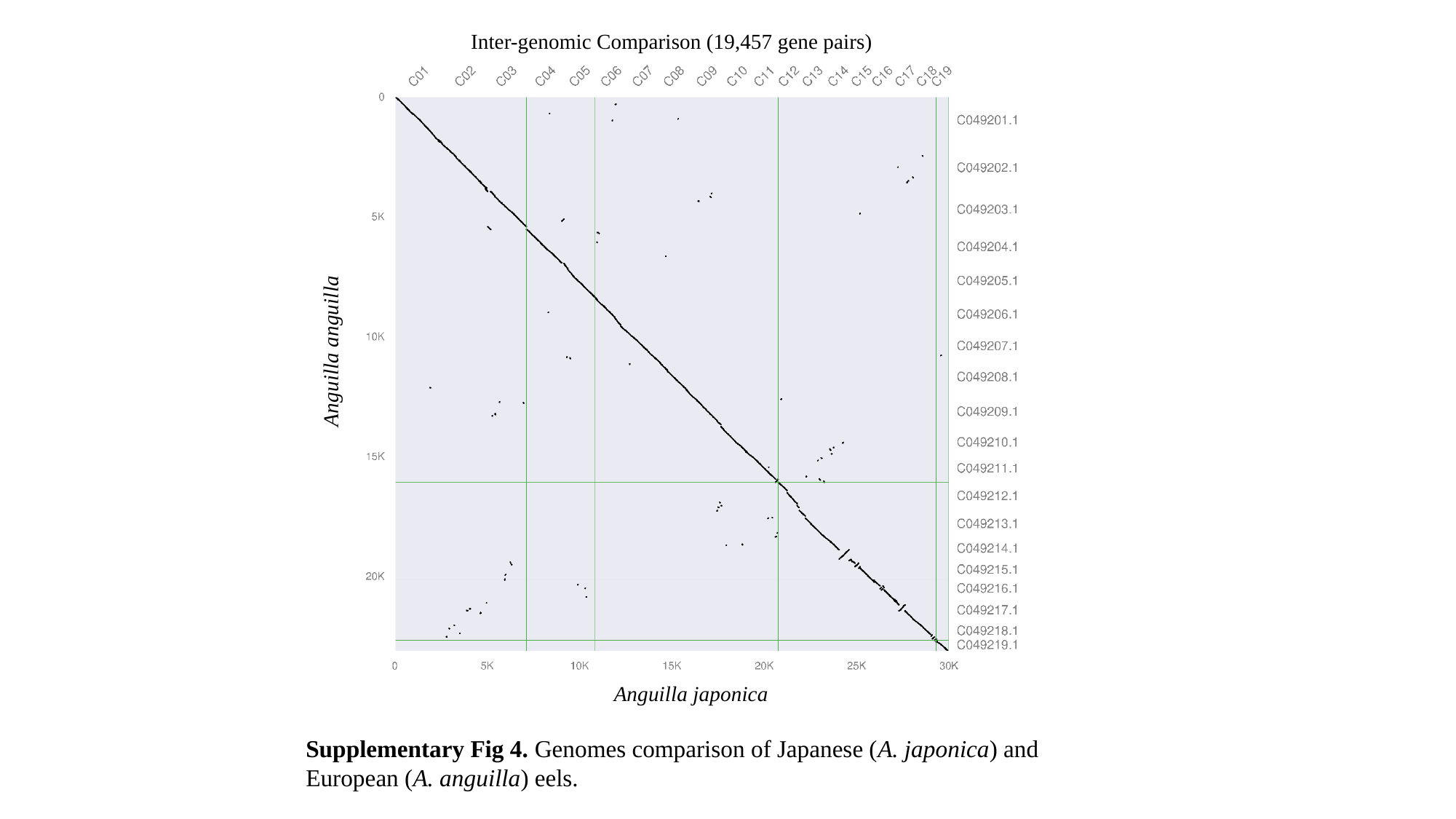

Inter-genomic Comparison (19,457 gene pairs)
Anguilla anguilla
Anguilla japonica
Supplementary Fig 4. Genomes comparison of Japanese (A. japonica) and European (A. anguilla) eels.

### Slide 5
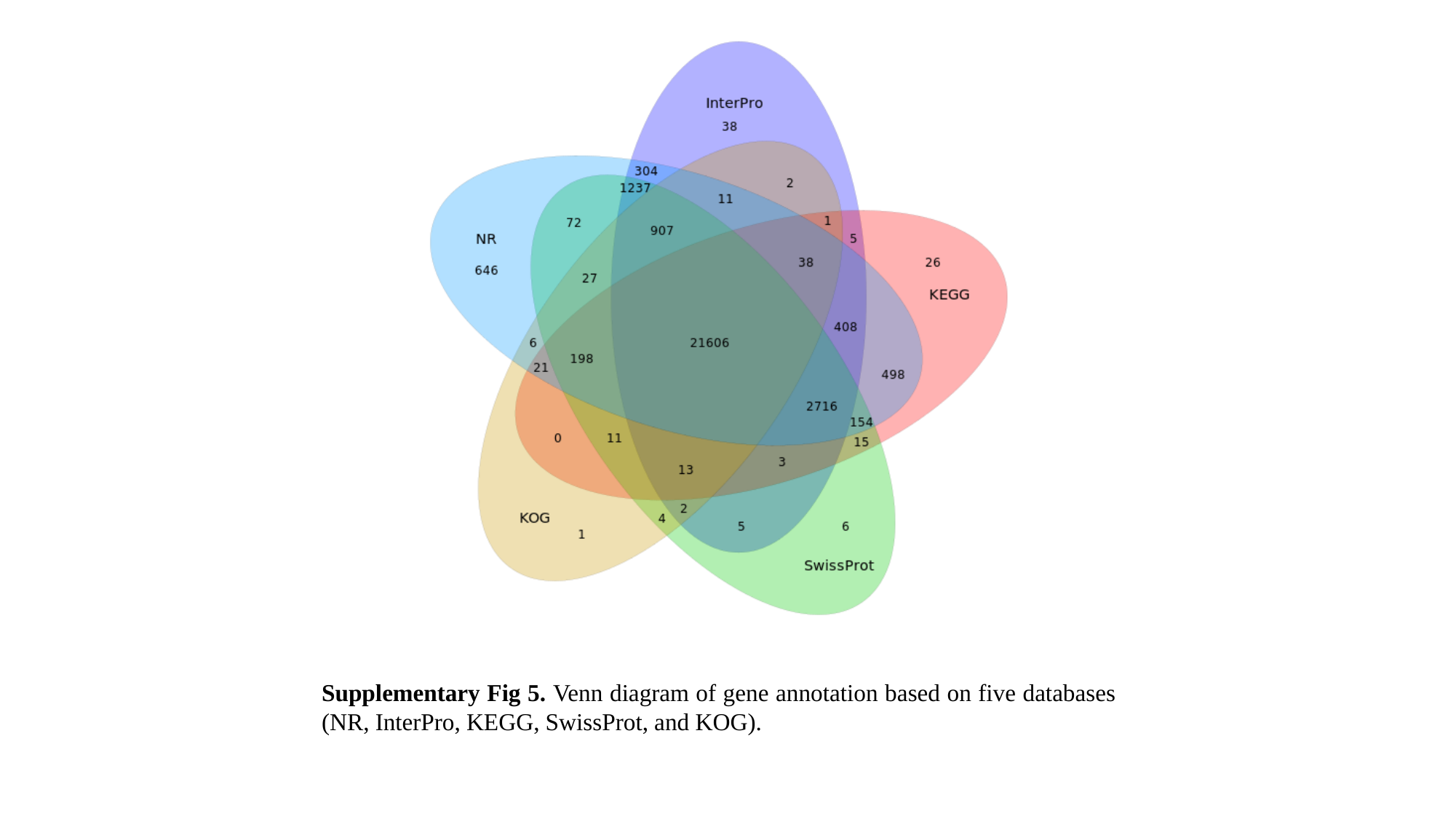

Supplementary Fig 5. Venn diagram of gene annotation based on five databases (NR, InterPro, KEGG, SwissProt, and KOG).

### Slide 6
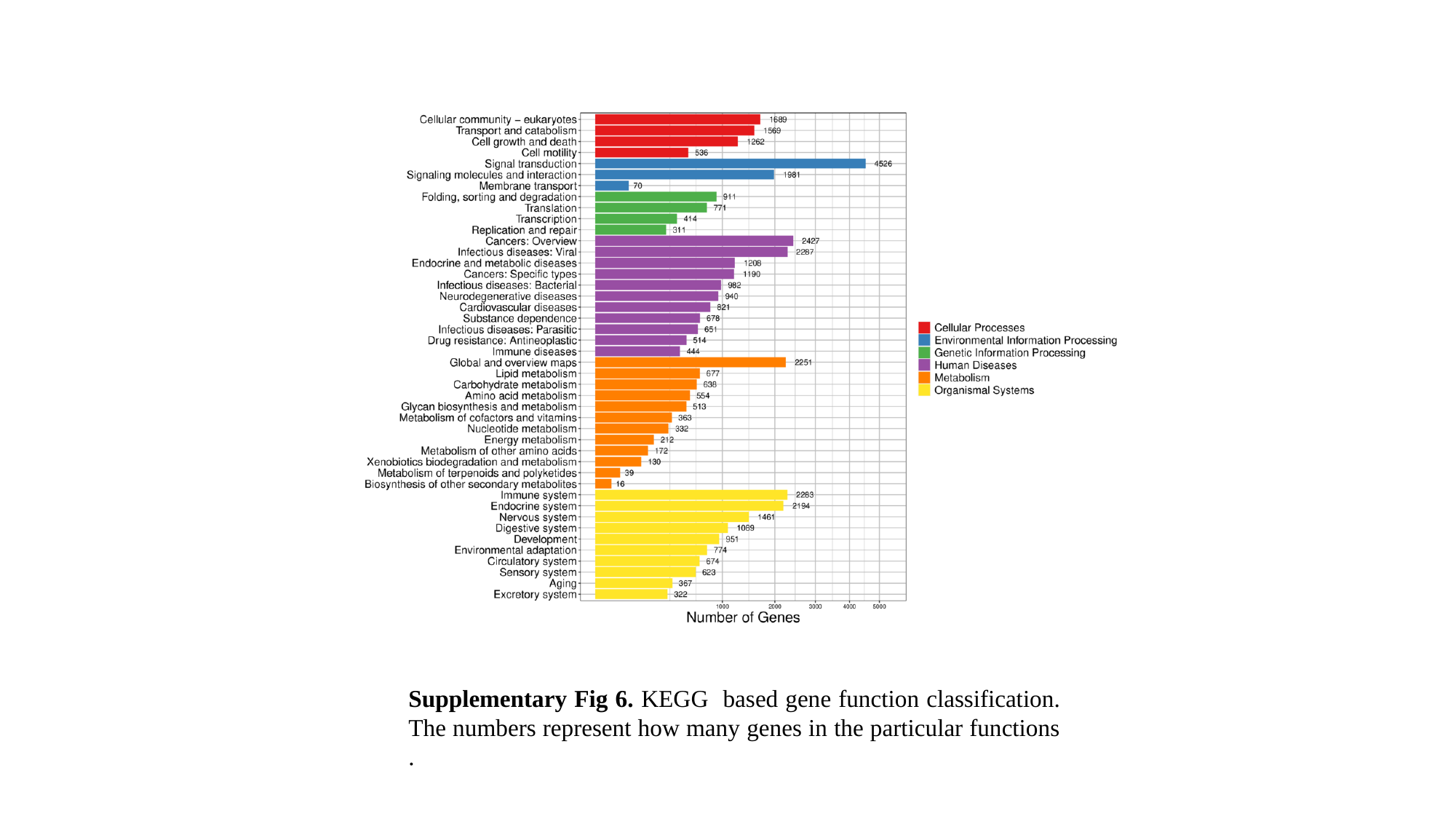

Supplementary Fig 6. KEGG based gene function classification. The numbers represent how many genes in the particular functions .

### Slide 7
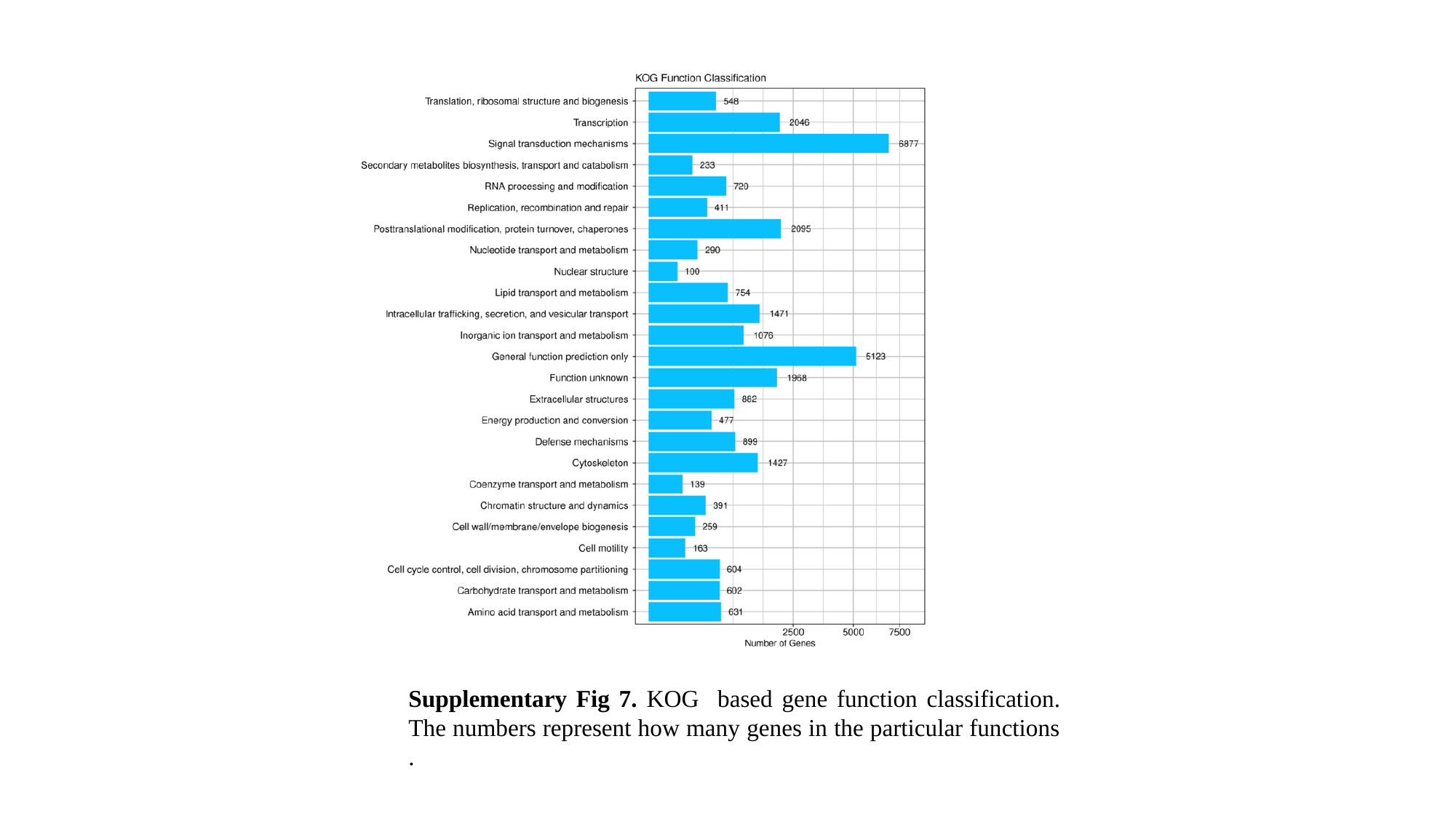

Supplementary Fig 7. KOG based gene function classification. The numbers represent how many genes in the particular functions .

### Slide 8
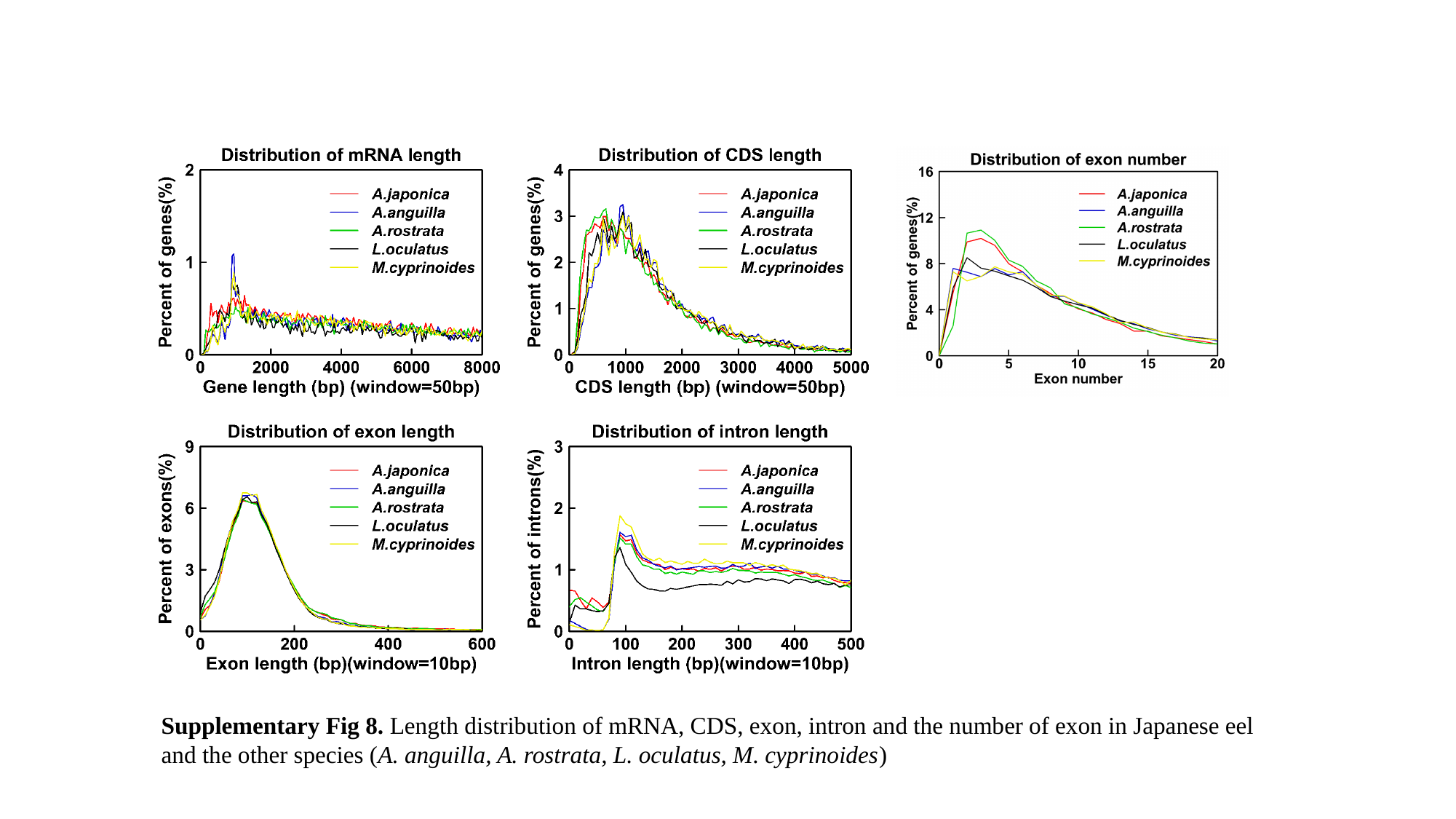

Supplementary Fig 8. Length distribution of mRNA, CDS, exon, intron and the number of exon in Japanese eel and the other species (A. anguilla, A. rostrata, L. oculatus, M. cyprinoides)

### Slide 9
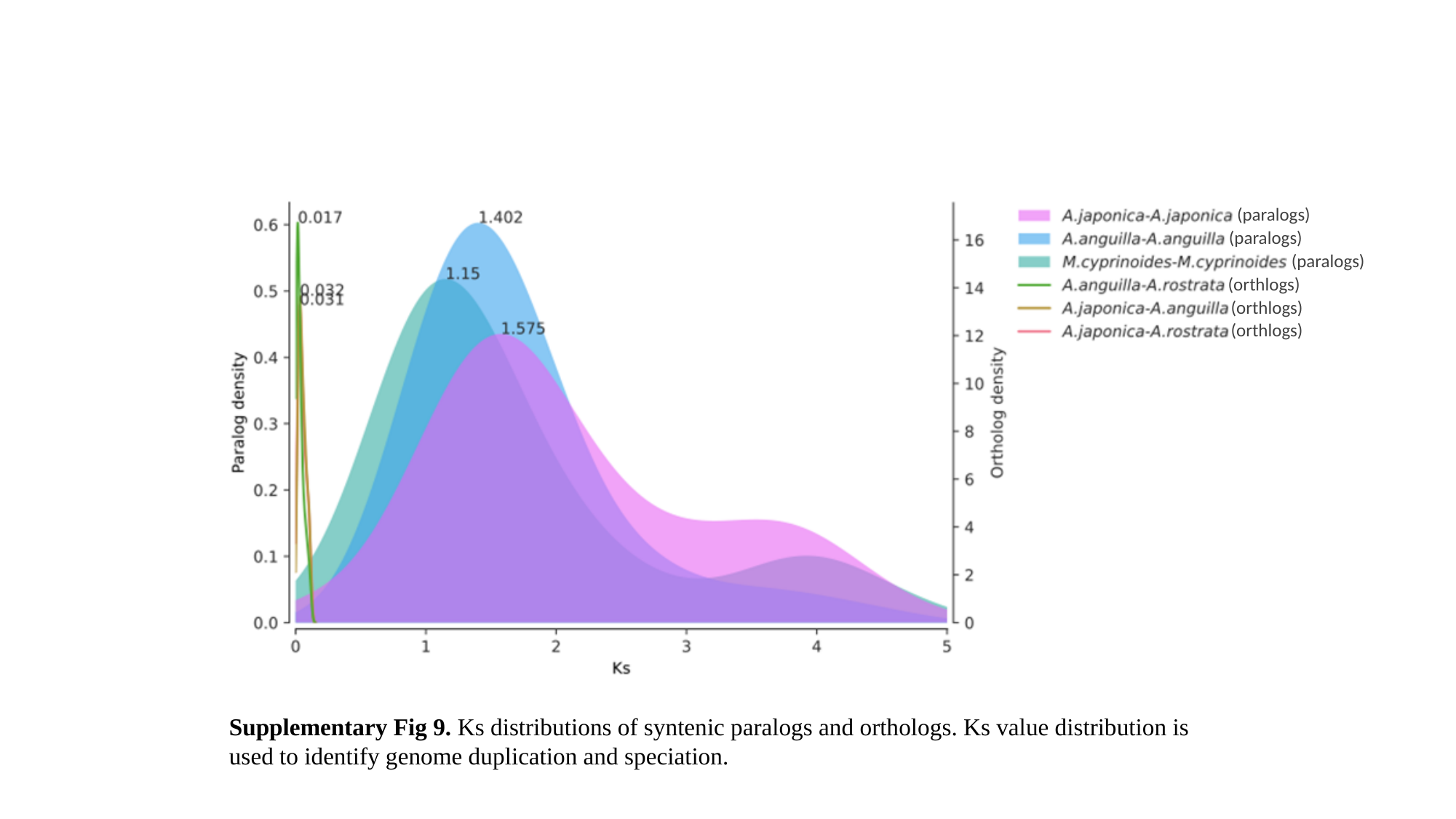

(paralogs)
(paralogs)
(paralogs)
(orthlogs)
(orthlogs)
(orthlogs)
Supplementary Fig 9. Ks distributions of syntenic paralogs and orthologs. Ks value distribution is used to identify genome duplication and speciation.

### Slide 10
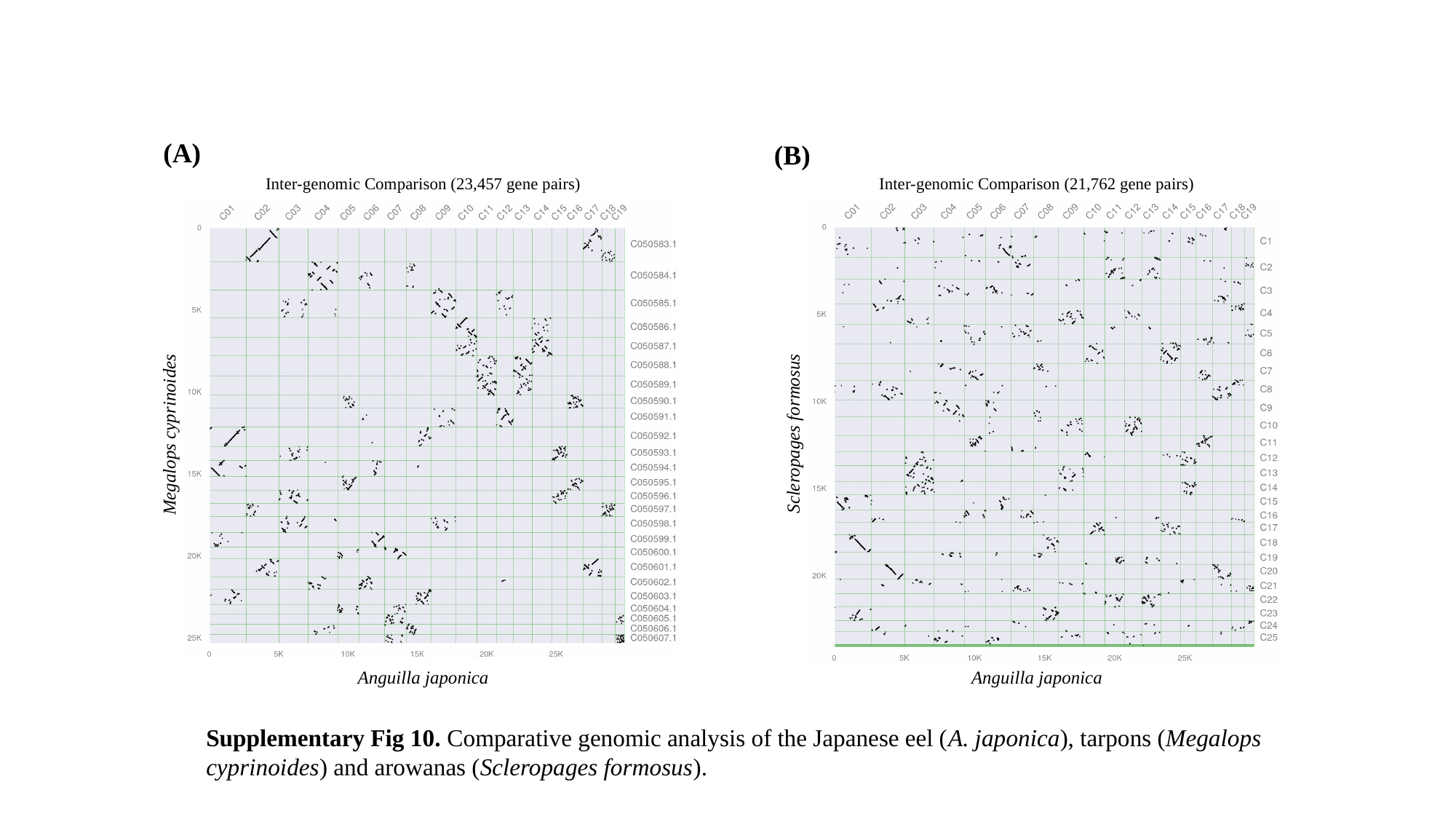

(A)
(B)
Inter-genomic Comparison (23,457 gene pairs)
Megalops cyprinoides
Anguilla japonica
Inter-genomic Comparison (21,762 gene pairs)
Scleropages formosus
Anguilla japonica
Supplementary Fig 10. Comparative genomic analysis of the Japanese eel (A. japonica), tarpons (Megalops cyprinoides) and arowanas (Scleropages formosus).
