## Supplementary material for "A Chromosome-level Assembly of the Japanese Eel Genome, Insights into Gene Duplication and Chromosomal Reorganization": Table 1

Statistics of *Anguilla japonica* genome assembly and annotation

| Assembly feature | *Anguilla japonica* |
| --- | --- |
| Genome size, Gb | 1.028 |
| No. of contigs | 811 |
| Contig N50, Mbp | 21.48 |
| Contig N90, Kbp | 716.98 |
| Longest contig, Mbp | 57.08 |
| No. of scaffolds | 86 |
| Scaffold N50, Mbp | 58.71 |
| Scaffold N90, Mbp | 38.29 |
| Longest scaffold, Mbp | 94.29 |
| Repeat portion of assembly, % | 30.48 |
| No. of genes | 29,982 |
| GC% | 44 |
| Genes average length, bp | 10265.73 |
| Average exons per gene | 9 |
